# α_3_β_4_* Nicotinic acetylcholine receptors strongly modulate the excitability of VIP neurons in the mouse inferior colliculus

**DOI:** 10.1101/2021.04.13.439708

**Authors:** Luis M. Rivera-Perez, Julia T. Kwapiszewski, Michael T. Roberts

## Abstract

The inferior colliculus (IC), the midbrain hub of the central auditory system, receives extensive cholinergic input from the pontomesencephalic tegmentum. Activation of nicotinic acetylcholine receptors (nAChRs) in the IC can alter acoustic processing and enhance auditory task performance. However, how nAChRs affect the excitability of specific classes of IC neurons remains unknown. Recently, we identified vasoactive intestinal peptide (VIP) neurons as a distinct class of glutamatergic principal neurons in the IC. Here, in experiments using male and female mice, we show that cholinergic terminals are routinely located adjacent to the somas and dendrites of VIP neurons. Using whole-cell electrophysiology in brain slices, we found that acetylcholine drives surprisingly strong and long-lasting excitation and inward currents in VIP neurons. This excitation was unaffected by the muscarinic receptor antagonist atropine. Application of nAChR antagonists revealed that acetylcholine excites VIP neurons mainly via activation of α_3_β_4_* nAChRs, a nAChR subtype that is rare in the brain. Furthermore, we show that cholinergic excitation is intrinsic to VIP neurons and does not require activation of presynaptic inputs. Lastly, we found that low frequency trains of acetylcholine puffs elicited temporal summation in VIP neurons, suggesting that in vivo-like patterns of cholinergic input can reshape activity for prolonged periods. These results reveal the first cellular mechanisms of nAChR regulation in the IC, identify a functional role for α_3_β_4_* nAChRs in the auditory system, and suggest that cholinergic input can potently influence auditory processing by increasing excitability in VIP neurons and their postsynaptic targets.

**Key points summary:**

- The inferior colliculus (IC), the midbrain hub of the central auditory system, receives extensive cholinergic input and expresses a variety of nicotinic acetylcholine receptor (nAChR) subunits.
- In vivo activation of nAChRs alters the input-output functions of IC neurons and influences performance in auditory tasks. However, how nAChR activation affects the excitability of specific IC neuron classes remains unknown.
- Here we show in mice that cholinergic terminals are located adjacent to the somas and dendrites of VIP neurons, a class of IC principal neurons.
- We find that acetylcholine elicits surprisingly strong, long-lasting excitation of VIP neurons and this is mediated mainly through activation of α_3_β_4_* nAChRs, a subtype that is rare in the brain.
- Our data identify a role for α_3_β_4_* nAChRs in the central auditory pathway and reveal a mechanism by which cholinergic input can influence auditory processing in the IC and the postsynaptic targets of VIP neurons.

## Introduction

Growing evidence indicates that cholinergic signaling through nicotinic acetylcholine receptors (nAChRs) critically shapes sound processing in the central auditory system (Goyer *et al*., 2016; Askew *et al*., 2017; Felix *et al*., 2019; Zhang *et al*., 2021). The inferior colliculus (IC), the midbrain hub of the central auditory system, receives extensive input from cholinergic neurons in the pontomesencephalic tegmentum (PMT; Motts & Schofield, 2009), and expresses several nAChR subunits, including α_3_, α_4_, α_7_, β_2_, β_3_, and β_4_ (Clarke *et al*., 1985; Wada *et al*., 1989; Morley & Happe, 2000; Whiteaker *et al*., 2002; Salas *et al*., 2003; Gahring *et al*., 2004; Happe & Morley, 2004; Bieszczad *et al*., 2012; Sottile *et al*., 2017). Because activity in the PMT is influenced by the sleep-wake cycle, attention, rewards, and sensory novelty, it is hypothesized that PMT neurons regulate auditory processing in the IC as a function of behavioral state (Jones, 1991; Kozak *et al*., 2005; Schofield *et al*., 2011; Boucetta *et al*., 2014). Consistent with this, in vivo studies have shown that nicotinic drugs alter the gain of input-output functions in IC neurons (Farley *et al*., 1983; Habbicht & Vater, 1996), and human psychophysics studies indicate that nicotine improves performance in auditory attention and discrimination tasks (Knott *et al*., 2012; Smucny *et al*., 2016; Pham *et al*., 2020), an effect partly attributable to alterations in the IC (Askew *et al*., 2017). In addition, temporal coding of auditory stimuli is degraded in the IC of α_7_ knockout mice (Felix *et al*., 2019). However, despite the importance of nAChRs to auditory processing, the cellular mechanisms by which nAChRs influence the excitability IC neurons remain largely unknown.

This gap in knowledge has eluded the field mostly due to the complexity of the neuronal populations in the IC, where it has proven difficult to identify and study specific neuron classes using conventional approaches. We recently overcame this obstacle, identifying vasoactive intestinal peptide (VIP) neurons as the first molecularly identifiable neuron class in the IC (Goyer et al., 2019). VIP neurons are found throughout the major IC subdivisions, they are glutamatergic, and they have a stellate morphology with spiny dendrites that, within the central nucleus of the IC (ICc), typically extend across two or more isofrequency laminae. VIP neurons project to several auditory regions, including the auditory thalamus, superior olivary complex, and the contralateral IC, and they receive input from the dorsal cochlear nucleus, the contralateral IC, and likely from other sources. By using the VIP-IRES-Cre mouse model, we can selectively target VIP neurons for electrophysiological and anatomical experiments. Thus, we are in a position for the first time to determine the cellular mechanisms of cholinergic signaling in a defined class of IC neurons.

Here, we hypothesized that the excitability of VIP neurons in the IC is modulated by cholinergic signaling. Using immunofluorescence, we showed that cholinergic terminals are frequently located in close proximity to VIP neurons, suggesting that VIP neurons receive direct cholinergic input. We then found that brief applications of ACh elicited surprisingly long periods of depolarization and spiking in VIP neurons. These responses were not affected by atropine, a muscarinic acetylcholine receptor (mAChR) antagonist, but were largely blocked by mecamylamine, an antagonist partially selective for β_4_-containing receptors, and by SR16584, an antagonist selective for α_3_β_4_* receptors (* *indicates that the identity of the fifth subunit in the receptor pentamer is unknown*). Consistent with this, voltage clamp recordings showed that ACh puffs led to prolonged inward currents that were largely blocked by mecamylamine. Moreover, cholinergic responses were resistant to manipulations affecting synaptic transmission, indicating that the nAChRs mediating these responses are expressed by VIP neurons.

Finally, we showed that 10 Hz trains of lower concentration ACh puffs elicited temporal summation in VIP neurons, suggesting that the in vivo firing patterns of cholinergic PMT neurons are likely to drive prolonged excitation of VIP neurons. We thus provide the first evidence that α_3_β_4_* nAChRs, a subtype with limited distribution in the brain, elicit direct and potent excitation of IC VIP neurons. Combined, our data reveal that cholinergic modulation exerts a surprisingly potent and long-lasting increase in the excitability of an important class of IC principal neurons.

## Materials and Methods

### Animals

All experiments were approved by the University of Michigan Institutional Animal Care and Use Committee and were in accordance with NIH guidelines for the care and use of laboratory animals. Animals were kept on a 12-hour day/night cycle with ad libitum access to food and water. VIP-IRES-Cre mice (*Vip*^*tm1(cre)Zjh*^/J, Jackson Laboratory, stock # 010908) (Taniguchi *et al*., 2011) were crossed with Ai14 reporter mice (B6.Cg-*Gt(ROSA)26Sor*^*tm14(CAG-tdTomato)Hze*^/J, Jackson Laboratory, stock #007914) (Madisen *et al*., 2010) to yield F1 offspring that expressed the fluorescent protein tdTomato in VIP neurons. Because mice on the C57BL/6J background undergo age-related hearing loss, experiments were restricted to mice aged P30 – P85, an age range where hearing loss should be minimal or not present (Zheng *et al*., 1999).

### Immunofluorescence

Mice aged P53 – P85 were deeply anesthetized with isoflurane and perfused transcardially with 0.1 M phosphate-buffered saline (PBS), pH 7.4, for 1 min and then with a 10% buffered formalin solution (Millipore Sigma, cat# HT501128) for 15 min. Brains were collected and post-fixed in the same formalin solution and cryoprotected overnight at 4°C in 0.1 M PBS containing 20% sucrose. Brains were cut into 40 µm sections on a vibratome. Sections were rinsed in 0.1 M PBS, and then treated with 10% normal donkey serum (Jackson ImmunoResearch Laboratories, West Grove, PA) and 0.3% TritonX-100 for 2 hr. Slices were incubated overnight at 4°C in rabbit anti-VAChT (3:500, Synaptic Systems, cat# 139103, RRID: AB_887864). This antibody was previously validated by Western blot and has been successfully used to identify cholinergic terminals in the cochlear nucleus and hippocampus (Gillet et al., 2018; Goyer et al., 2016; Zhang et al., 2019). The next day, sections were rinsed in 0.1 M PBS and incubated in Alexa Fluor 647-tagged donkey anti-rabbit IgG (1:500, ThermoFisher, cat# A-31573) for 2 hr at room temperature. Sections were then mounted on slides (SouthernBiotech, cat# SLD01-BX) and coverslipped using Fluoromount-G (SouthernBiotech, cat# 0100–01). Images were collected using a 1.40 NA 63x oil-immersion objective and 0.1 µm Z-steps on a Leica TCS SP8 laser scanning confocal microscope.

### Analysis of cholinergic terminals adjacent to VIP neurons

After immunofluorescence was performed, we used Neurolucida 360 software (MBF Bioscience) to reconstruct VIP neurons and assess the distribution of cholinergic terminals on reconstructed neurons. Terminals that were < 2 μm from the dendrites or soma of the reconstructed cell were counted as synapses onto that neuron.

### Brain slice preparation

Whole-cell patch-clamp recordings were performed in acutely prepared brain slices from VIP-IRES-Cre x Ai14 mice. Both males (n = 35) and females (n = 24) aged P30-P50 were used. No differences were observed between animals of different sexes. Mice were deeply anesthetized with isoflurane and then rapidly decapitated. The brain was removed, and a tissue block containing the IC was dissected in 34°C ACSF containing the following (in mM): 125 NaCl, 12.5 glucose, 25 NaHCO_3_, 3 KCl, 1.25 NaH_2_PO_4_, 1.5 CaCl_2_ and 1 MgSO_4_, bubbled to a pH of 7.4 with 5% CO_2_ in 95% O_2_. Coronal sections of the IC (200 µm) were cut in 34°C ACSF with a vibrating microtome (VT1200S, Leica Biosystems) and incubated at 34°C for 30 min in ACSF bubbled with 5% CO_2_ in 95% O_2_. Slices were then incubated at room temperature for at least 30 min before being transferred to the recording chamber.

### Current-clamp electrophysiology

Slices were placed in a recording chamber under a fixed stage upright microscope (BX51WI, Olympus Life Sciences) and were constantly perfused with 34 °C ACSF at ∼2 ml/min. All recordings were conducted near physiological temperature (34 °C). IC neurons were patched under visual control using epifluorescence and Dodt gradient-contrast imaging. Current-clamp recordings were performed with a BVC-700A patch clamp amplifier (Dagan Corporation). Data were low pass filtered at 10 kHz, sampled at 50 kHz with a National Instruments PCIe-6343 data acquisition board, and acquired using custom written algorithms in Igor Pro. Electrodes were pulled from borosilicate glass (outer diameter 1.5 mm, inner diameter 0.86 mm, Sutter Instrument) to a resistance of 3.5 – 5.0 MΩ using a P-1000 microelectrode puller (Sutter Instrument). The electrode internal solution contained (in mM): 115 Kgluconate, 7.73 KCl, 0.5 EGTA, 10 HEPES, 10 Na_2_ phosphocreatine, 4 MgATP, 0.3 NaGTP, supplemented with 0.1% biocytin (w/v), pH adjusted to 7.4 with KOH and osmolality to 290 mmol/kg with sucrose. Data were corrected for an 11 mV liquid junction potential.

To test the effect of ACh on the excitability of VIP neurons, acetylcholine chloride (Sigma cat # A6625), was freshly dissolved each day in a vehicle solution containing (in mM): 125 NaCl, 3 KCl, 12.5 Glucose and 3 HEPES. The solution was balanced to a pH of 7.40 with NaOH. The working concentration of ACh was 1 mM unless stated otherwise. To apply ACh puffs on brain slices, ACh solution was placed in pipettes pulled from borosilicate glass (outer diameter 1.5 mm, inner diameter 0.86 mm, Sutter Instrument) with a resistance of 3.5 – 5.0 MΩ using a P-1000 microelectrode puller (Sutter Instrument) connected to a pressure ejection system built based on the OpenSpritzer design (Forman *et al*., 2017). The puffer pipette was placed ∼ 20 um from the soma of the patched cell, and five 10 ms puff applications at 10 psi and 1 minute apart were presented per condition. To isolate the receptors mediating the effects of ACh on VIP neurons, we bath applied the following drugs individually or in combination: 1 μM atropine (mAChR antagonist, Sigma), 5 μM mecamylamine (Mec, relatively non-selective antagonist with higher affinity for β_4_ containing receptors, Sigma), 10 µM DHβE (α_4_β_2_* nAChR antagonist, Tocris), 50 µM SR16584 (α_3_β_4_* nAChR antagonist, Tocris), and 5 nM methyllycaconitine (MLA, α_7_ nAChR antagonist, Sigma). All drugs were washed-in for 10 minutes before testing how the drugs affected the responses of the recorded neurons to ACh puffs. In one experiment, antagonists for GABA_A_, glycine, AMPA, and NMDA receptors were bath applied to isolate direct effects of ACh on VIP neurons from possible ACh-induced changes in release from terminals synapsing onto VIP neurons. The following drug concentrations were used: 5 µM SR95531 (gabazine, GABA_A_ receptor antagonist, Hello Bio), 1 µM strychnine hydrochloride (glycine receptor antagonist, Millipore Sigma), 10 µM NBQX disodium salt (AMPA receptor antagonist, Hello Bio), 50 µM D-AP5 (NMDA receptor antagonist, Hello Bio). All drugs were washed-in for 10 minutes before testing how the drugs affected the responses of the recorded neurons to ACh puffs. Except for when the effects of atropine alone were directly tested, 1 μM atropine was included in the ACSF under all conditions.

### Effect of repeated ACh applications

ACh puffs were applied as described above except at lower concentrations (30 µM and 100 µM). Trials containing trains of 1, 2, 3, 5 or 10 puffs at 10 Hz were delivered with a 1-minute intertrial period. Five trials were presented per condition.

### Voltage-clamp recordings of nAChR currents

For voltage-clamp experiments, the recording setup was the same as above except that recordings were performed using an Axopatch 200A amplifier. During the recordings, the series resistance compensation was performed using 90% prediction and 90% correction. The series resistance of the electrode was never greater than 10 MΩ. The electrode internal solution contained (in mM): 115 CsOH, 115 D-gluconic acid, 7.76 CsCl, 0.5 EGTA, 10 HEPES, 10 Na_2_ phosphocreatine, 4 MgATP, 0.3 NaGTP, supplemented with 0.1% biocytin (w/v), pH adjusted to 7.4 with CsOH and osmolality to 290 mmol/kg with sucrose. As detailed above, the ACh puffer was placed approximately 20 µm from the soma of the patched cell, and five 10 ms puff applications at 10 psi were presented per condition, waiting 1 minute between puffs. Receptor antagonists were applied as described above for the current clamp experiments. 1 μM atropine was included in the ACSF under all conditions. Voltage-clamp holding potentials were not corrected for the liquid junction potential.

### Analysis of electrophysiological recordings

Data analysis and statistical tests were performed using custom written algorithms in Igor Pro (Wavemetrics) and MATLAB (MathWorks). Statistical tests included paired t-tests, repeated measure ANOVAs with Tukey’s post hoc, and linear regression analysis, as indicated in the figure legends. Data are presented as mean ± SD and were considered significant when p < 0.05.

## Results

### Cholinergic synapses are found adjacent to the somas and dendrites of VIP neurons

The IC receives cholinergic input from the two nuclei that comprise the PMT: the pedunculopontine tegmental nucleus and the laterodorsal tegmental nucleus. Together, these nuclei distribute cholinergic axons and synapses throughout the IC, contacting both GABAergic and glutamatergic neurons (Motts & Schofield, 2009; Schofield *et al*., 2011; Noftz *et al*., 2020). However, the specific neuronal populations that cholinergic terminals synapse onto in the IC remain unclear. To test whether VIP neurons receive cholinergic input, we performed immunofluorescence on brain slices from VIP-IRES-Cre x Ai14 mice, in which VIP neurons express the fluorescent protein tdTomato, using an antibody against the vesicular acetylcholine transporter (VAChT). High resolution images were collected using a laser-scanning confocal microscope with a 1.40 NA 63x oil-immersion objective and 0.1 µm Z-steps. Analysis of these images showed that VAChT^+^ boutons and terminals were routinely located < 2 µm from the somas, dendrites, or both of VIP neurons (**Figure 1**). Similar results were observed in IC sections from five mice. These results suggest that VIP neurons receive cholinergic input (Rees *et al*., 2017). We therefore hypothesized that cholinergic signaling modulates the excitability of VIP neurons.

**Figure 1:**
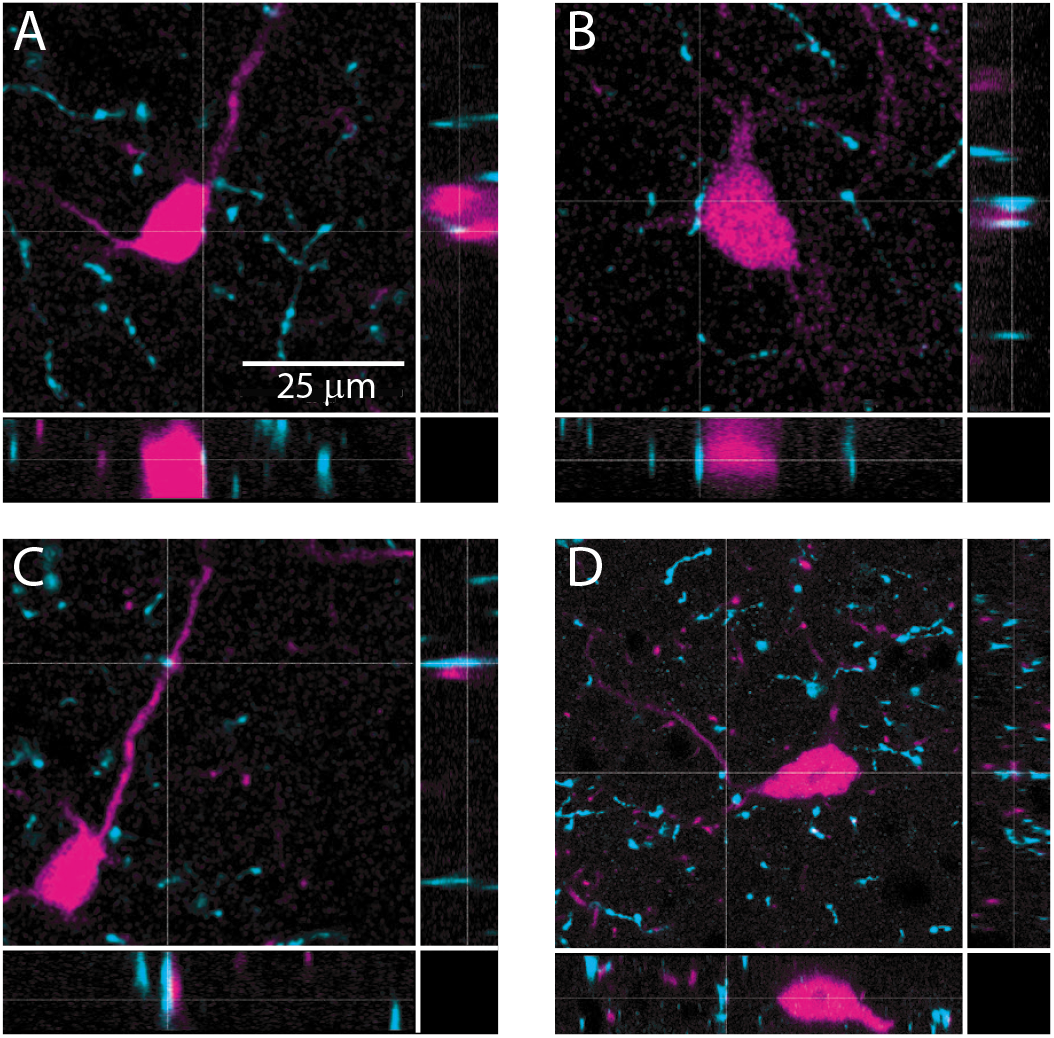
Cholinergic terminals are routinely found in close proximity to VIP neuron somas and dendrites. Confocal images from IC sections show examples of cholinergic terminals (identified by dashed crosshairs) labeled by anti-VAChT (cyan) located < 2 µm from the somas (top row) or dendrites (bottom row) of VIP neurons (magenta). The three panels in each image provide a top view and two side views centered on the cholinergic terminal identified by the crosshairs. Images are from 3 mice.

### Brief puffs of ACh drive prolonged firing in VIP neurons via non-α_7_ nAChRs

To test whether acetylcholine alters the excitability of VIP neurons, we targeted current clamp recordings to fluorescent VIP neurons in acute IC slices from VIP-IRES-Cre x Ai14 mice and used a puffer pipette to provide brief puffs of ACh near the recorded cell. We found that 10 ms puffs of 1 mM ACh delivered approximately 20 µm from the VIP cell soma drove depolarization and firing in 97 out of 107 VIP neurons. These effects were surprisingly strong and long-lasting, suggesting that cholinergic signaling can potently increase the excitability of VIP neurons (**Figure 2**).

**Figure 2:**
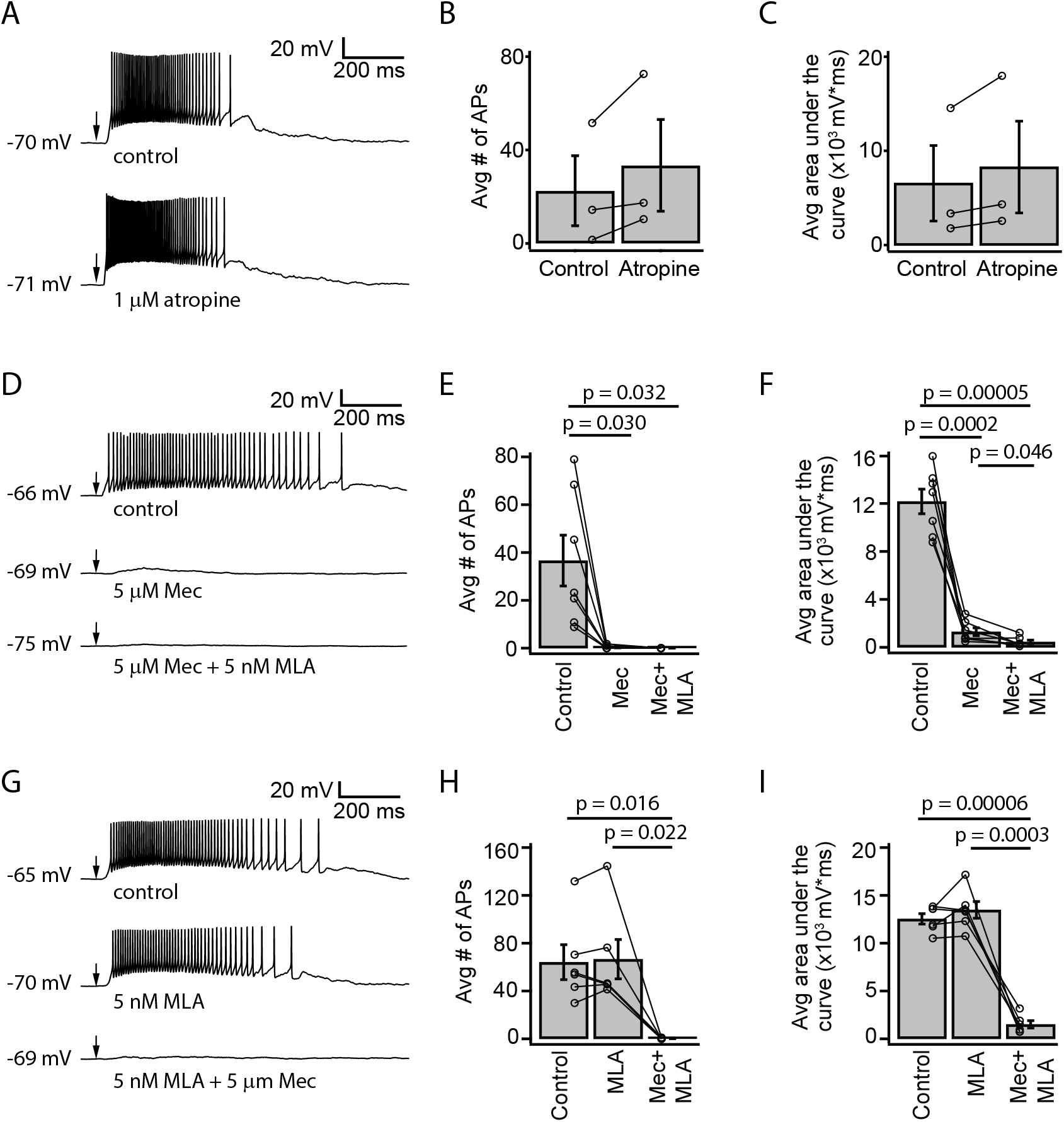
ACh-induced depolarization of VIP neurons is mediated by non-α7 nAChRs. *A*, A 10 ms puff of 1 µM ACh elicited a long-lasting depolarization and spiking in a representative VIP neuron under control conditions (top). 1 µM atropine did not alter the ACh response (bottom). *B,C*, Atropine did not change the average number of action potentials elicited by ACh or the average area under the median-filtered curve. (n = 3). Note that in all remaining experiments, 1 µM atropine was always present in the ACSF. *D*, Example traces show ACh-elicited depolarization in a VIP neuron in control ACSF (top), 5 µM Mec (middle), and 5 µM Mec + 5 nM MLA (bottom). *E*, Mec reduced the average number of action potentials elicited by ACh puffs from 36.6 ± 28.2 to 0.4 ± 0.7. (repeated measures ANOVA with Tukey’s post hoc, *F*_*2,12*_ = 11.99; n = 7; control vs Mec, p = 0.030; control vs Mec + MLA, p = 0.032; Mec vs Mec + MLA, p = 0.37). *F*, The total depolarization elicited by ACh puffs, measured as the area under the median filtered curve, was significantly decreased by Mec and further decreased by MLA (repeated measures ANOVA with Tukey’s post hoc, *F*_*2,12*_ = 111.6; n = 7; control vs Mec, p = 0.0002; control vs Mec + MLA, p = 4.7 × 10^−5^; Mec vs Mec + MLA, p = 0.046). *G*, When MLA was applied before Mec, MLA alone did not have a clear effect on ACh responses. From top to bottom: control, 5 nM MLA, and 5 µM Mec + 5 nM MLA. *H*, The average number of action potentials elicited by ACh was not significantly affected by MLA, while subsequent application of Mec + MLA eliminated firing (repeated measures ANOVA with Tukey’s post hoc, *F*_*2,10*_ = 17.23; n = 6; control vs MLA, p = 0.80; control vs MLA + Mec, p = 0.016; MLA vs MLA + Mec, p = 0.022). *I*, MLA alone did not significantly alter the depolarization elicited by ACh, while subsequent application of Mec + MLA caused an 89% reduction in the average depolarization (repeated measures ANOVA with Tukey’s post hoc, *F* _*2,10*_ = 120.4; n = 6; control vs MLA, p = 0.39; control vs MLA + Mec, p = 5.6 × 10^−5^; MLA vs MLA + Mec, p = 0.0003). In *A, D, G*, arrows indicate the time of the ACh puffs, and voltages indicate resting membrane potential.

Since ACh depolarized VIP neurons for up to 1 second, we first hypothesized that this effect was mediated by a slow metabotropic mechanism involving muscarinic acetylcholine receptors (mAChRs). However, we found that ACh-mediated excitation of VIP neurons was not altered by 1 µM atropine, a mAChR antagonist, indicating that mAChRs are not involved in this phenomenon (**Figure 2A-C**). For the remainder of this study, all recordings were conducted in the presence of 1 µM atropine, allowing us to isolate effects on nAChRs.

Next, we used mecamylamine (Mec), a broad-spectrum nAChR antagonist partially selective for β4-containing nAChRs (Papke *et al*., 2008, 2010), and methyllycaconitine (MLA), an antagonist selective for α_7_ nAChRs, to assess the contributions of nAChRs to the cholinergic excitation of VIP neurons. We found that bath application of 5 µM Mec nearly abolished the firing and strongly reduced the depolarization elicited by ACh puffs on VIP neurons. When both 5 µM Mec and 5 nM MLA were applied, the remaining depolarization was nearly eliminated (**Figure 2D-F**). When 5 nM MLA was applied first, ACh-elicited firing and depolarization in VIP neurons were not significantly altered. Subsequent addition of 5 µM Mec to the bath abolished the ACh effect (**Figures 2G-I**). Combined, these results suggest that cholinergic modulation of VIP neurons is predominately driven by non-α_7_, Mec-sensitive nAChRs.

### Brief ACh puffs elicit a long-lasting inward current in VIP neurons

nAChRs are commonly associated with fast, short-lasting depolarizations, but our data suggest that activation of nAChRs elicits prolonged depolarization in VIP neurons. To analyze the currents generated by activation of nAChRs in VIP neurons, we used voltage-clamp recordings with the holding potential at −60 mV. We found that a 10 ms puff of 1 mM ACh elicited an inward current in VIP neurons that lasted hundreds of milliseconds (mean decay τ = 438 ± 173 ms, mean ± SD, based on exponential fit; n = 5 neurons; **Figure 3A**). The peak of the ACh-evoked inward current was −329 ± 154 pA, and the 10 – 90% rise time was 89 ± 31 ms (mean ± SD, n = 5 neurons). Furthermore, similar to the depolarizations observed in our current-clamp experiments, 5 µM Mec abolished most of the current elicited by ACh, and the combination of 5 µM Mec and 5 nM MLA abolished the elicited current completely (**Figure 3B**). Therefore, our results suggest that the nAChRs mediating the effect of ACh on VIP neurons remain activated for extended periods, presumably due to slow kinetics and/or limited desensitization. Since α_7_ nAChRs have fast kinetics and rapid desensitization (Castro & Albuquerque, 1993; Anand *et al*., 1998; Papke *et al*., 2000; Mike *et al*., 2000), both the pharmacology and kinetics of the inward currents observed here are consistent with a mechanism mediated by non-α_7_ nAChRs.

**Figure 3:**
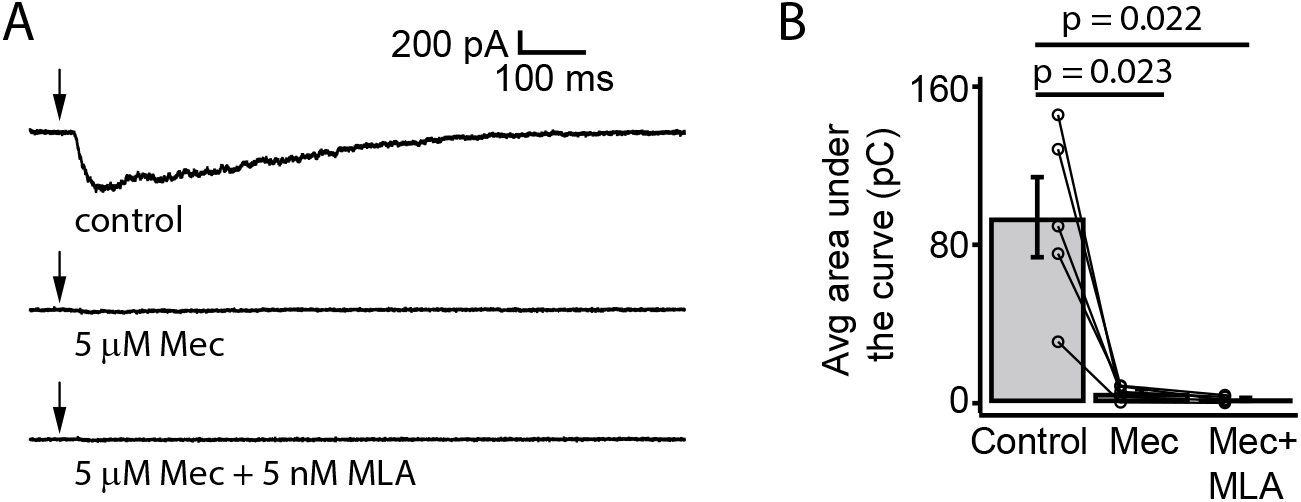
Brief ACh puffs elicit long-lasting inward currents in VIP neurons. *A*, Example traces of currents elicited by ACh puffs in control ACSF (top), 5 µM Mec (middle), and 5 µM Mec + 5 nM MLA (bottom). Arrows indicate the time of the ACh puffs. *B*, The total charge flux elicited by ACh, measured by the average area under the curve, significantly decreased after bath application of Mec. Subsequent application of MLA did not cause a further reduction (repeated measures ANOVA with Tukey’s post hoc, *F*_*2,8*_ = 20.5; n = 5; control vs Mec, p = 0.023; control vs Mec + MLA, p = 0.022; Mec vs Mec + MLA, p = 0.098).

### ACh-driven firing in VIP neurons does not require activation of presynaptic nAChRs

Many glutamatergic and GABAergic neurons in the IC express nAChRs (Sottile *et al*., 2017). In addition, nAChRs are often located on presynaptic terminals where their activation can directly promote neurotransmitter release (Dani & Bertrand, 2007). Therefore, it is possible that the ACh-elicited excitation of VIP neurons requires activation of an intermediate population of neurons or terminals that in turn excite VIP neurons through the release of a different, non-cholinergic neurotransmitter. We therefore tested if cholinergic modulation of VIP neurons requires activation of receptors for glutamate, GABA, and/or glycine, the main neurotransmitters in the IC. By using pharmacology to block these receptors (10 µM NBQX to block AMPA receptors, 50 µM D-APV to block NMDA receptors, 5 µM gabazine to block GABA_A_ receptors, and 1 µM strychnine to block glycine receptors), we isolated the effects of ACh puffs on VIP neurons from most other potential inputs. After bath application of the synaptic blockers, we observed that the depolarization elicited by ACh persisted and was not significantly altered (**Figure 4A**).

**Figure 4:**
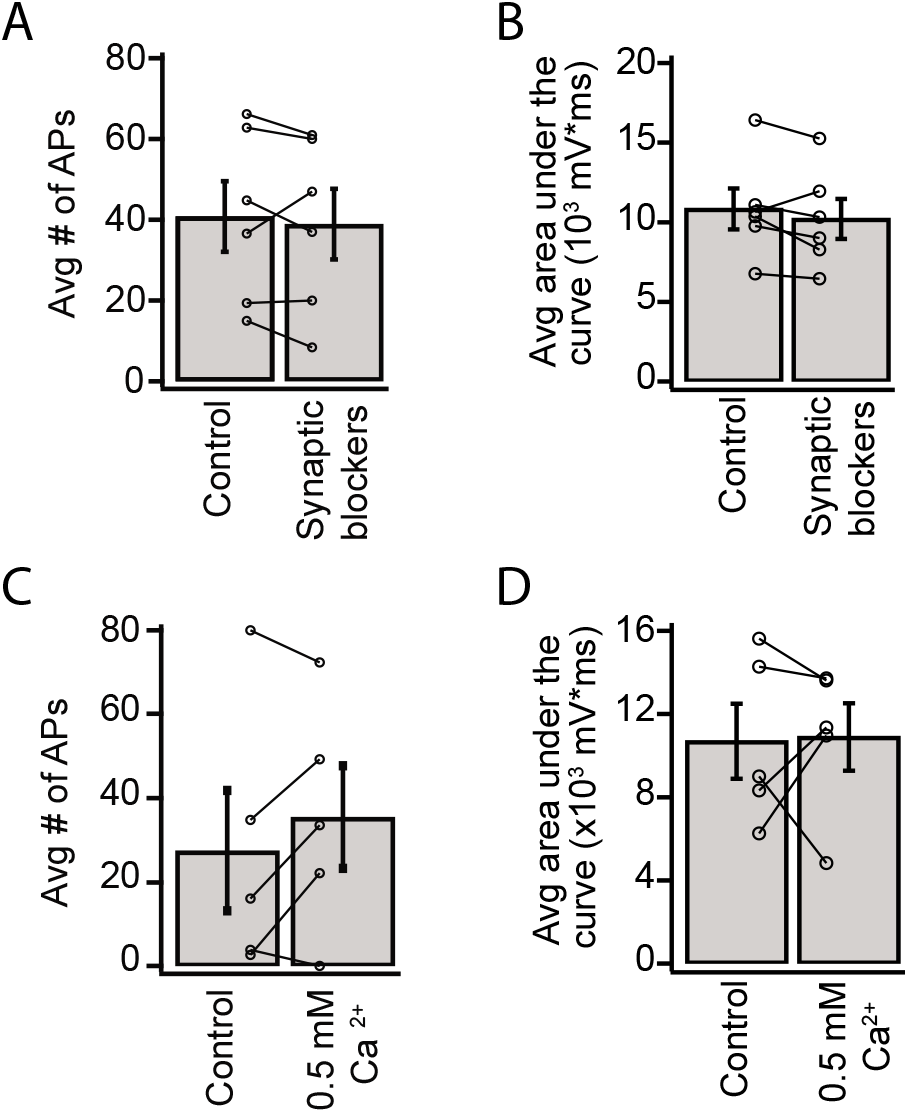
Depolarization of VIP neurons by ACh is consistent with an intrinsic mechanism and not presynaptic effects. *A,B*, The average number of action potentials (*A*) and depolarization (*B*) elicited by an ACh puff were not affected by bath application of antagonists against AMPA, NMDA, GABA_A_, and glycine receptors (10 µM NBQX, 50 µM D-AP5, 5 µM SR95537, and 1 µM strychnine; action potentials - paired t-test, *t*_*5*_ = 0.69; n = 6; p = 0.52; area under the curve – paired t-test, *t*_*5*_ = 1.37; n = 6; p = 0.23). *C,D*, The average number of action potentials (*C*) and depolarization (*D*) elicited by an ACh puff were unaffected by decreasing the ACSF Ca^2+^ concentration from 1.5 to 0.5 mM (action potentials – paired t-test, *t*_*4*_ = −1.4; n = 5; p = 0.23; area under the curve – paired t-test, *t*_*4*_ = −0.12; n = 5; p = 0.91).

Next, we globally reduced synaptic release probability by decreasing the concentration of Ca^2+^ in the ACSF from 1.5 mM to 0.5 mM. Since the relationship between release probability and extracellular Ca^2+^ is described by a power law (Dodge & Rahamimoff, 1967), this reduction in ACSF Ca^2+^ should dramatically decrease neurotransmitter release. We observed that decreasing extracellular Ca^2+^ did not significantly alter the depolarization elicited by ACh puffs on VIP neurons (**Figure 4B**). These results suggest that ACh acts on nAChRs present on VIP neurons themselves, and not via activation of presynaptic nAChRs.

### α_4_β_2_* nAChRs do not mediate the effect of ACh on VIP neurons

Our results thus far indicate that Mec-sensitive nAChRs mediate most of the effect of ACh on VIP neurons. However, Mec is a relatively broad-spectrum antagonist of non-homomeric nAChRs, with subtype selectivity depending on the concentration used (Papke *et al*., 2001, 2008, 2010). Since α_4_β_2_* nAChRs are widely expressed in the IC and are the most common subtype of nAChR found in the brain (Millar & Gotti, 2009), we performed current-clamp recordings to assess how DHβE, a selective antagonist for α_4_β_2_* nAChRs, affected the response of VIP neurons to ACh puffs. After bath-applying 10 µM DHβE for 10 minutes, our results showed that blocking α_4_β_2_* nAChRs did not significantly alter the spiking or depolarization elicited by ACh application (**Figure 5**). Therefore, our data suggest that cholinergic modulation of VIP neurons involves little or no contribution from α_4_β_2_* or α_7_ nAChRs, the most common nAChRs in the brain.

**Figure 5:**
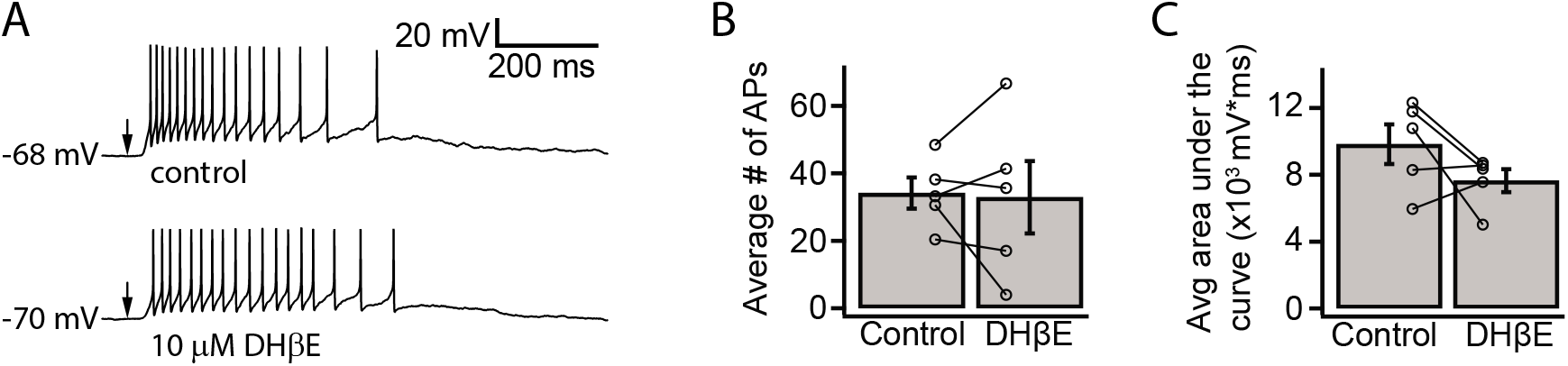
α_4_β_2_* nAChRs do not mediate the ACh-induced depolarization of VIP neurons. *A*, Example traces show that a spike train elicited by a 10 ms puff of 1 mM ACh (top) was not blocked by 10 µM DHβE (bottom), an α_4_β_2_* nAChR antagonist. Arrows indicate the time of the ACh puffs, and voltages indicate resting membrane potential. *B,C*, DHβE did not significantly affect the average number of action potentials (*B*) or the total depolarization (*C*) elicited by ACh puffs (action potentials – paired t-test, *t*_*4*_ = 0.17; n = 5; p = 0.88; area under the curve – paired t-test, *t*_*4*_ = 1.61; n = 5; p = 0.18).

### ACh-driven excitation of VIP neurons is mediated by α_3_β_4_* nAChRs

Although α_3_β_4_* nAChRs are relatively rare in the brain, previous studies indicate that α_3_ and β_4_ nAChR subunits are expressed in the IC (Wada *et al*., 1989; Marks *et al*., 2002, 2006; Whiteaker *et al*., 2002; Salas *et al*., 2003; Gahring *et al*., 2004). In addition, Mec strongly antagonizes α_3_β_4_* nAChRs at a concentration of 5 µM (Papke *et al*., 2008, 2010), which we used in our current-clamp and voltage-clamp recordings. We therefore hypothesized that α_3_β_4_* nAChRs mediate the excitatory effect of ACh on VIP neurons. To test this, we used SR16584, a selective α_3_β_4_* nAChR antagonist (Zaveri *et al*., 2010). Because SR16584 is dissolved in DMSO, we first established that a vehicle control (1:1000 DMSO:ACSF) did not affect the ability of 10 ms puffs of 1 mM ACh to excite VIP neurons (**Figure 6**). Next, we bath applied 50 µM SR16584 and found that it nearly abolished the spiking and strongly reduced the depolarization elicited by ACh (**Figure 6**), similar to our results with Mec applications. Furthermore, after only a 10-minute washout of SR16584, the excitatory effect of ACh partially recovered. These results demonstrate that the ACh-induced excitation of VIP neurons is mediated mainly by α_3_β_4_* nAChRs and provide the first evidence for a functional role of α_3_β_4_* nAChRs in the IC.

**Figure 6:**
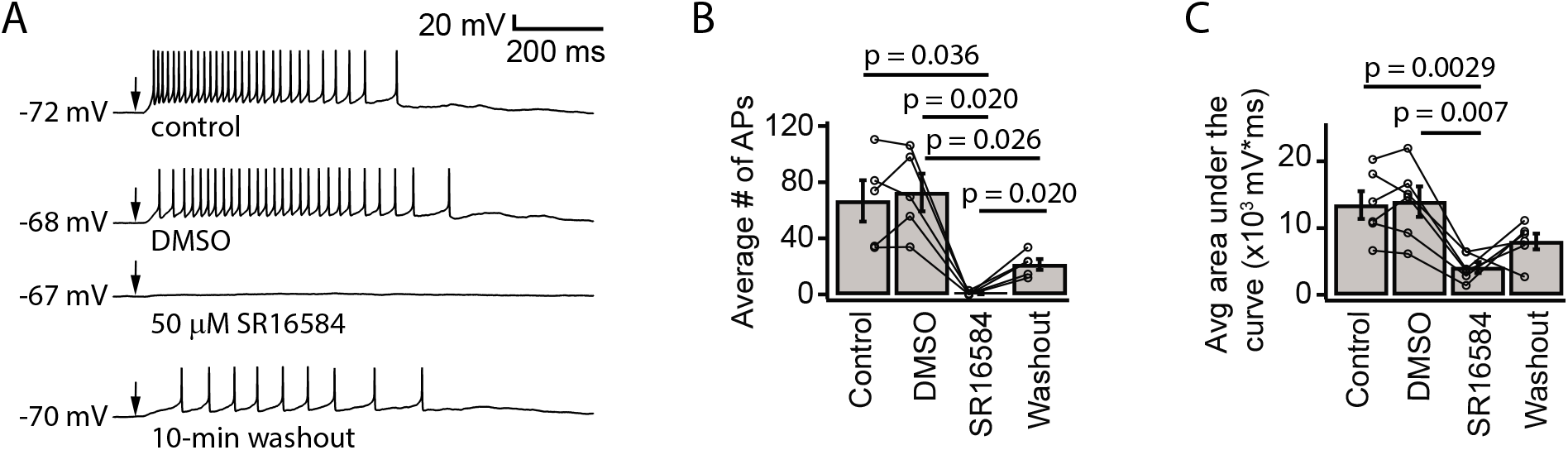
ACh-induced depolarization of VIP neurons is predominately mediated by α_3_β_4_* nAChRs. *A*, Example traces show that 50 µM SR16584, an α_3_β_4_* nAChR antagonist, blocked action potential firing and most of the depolarization elicited by 1 mM ACh puffs. Conditions from top to bottom: control, vehicle (1:1000 DMSO:ACSF), 50 µM SR16584, and 10-min washout (control ACSF). Arrows indicate the time of the ACh puffs, and voltages indicate resting membrane potential. *B*, The average number of action potentials elicited by ACh was reduced to 0.64 ± 1.22 after bath application of SR16584 (repeated measures ANOVA with Tukey’s post hoc, *F*_*3,12*_ = 20.3; n = 5; control vs DMSO, p = 0.83; control vs SR16584, p = 0.036; control vs washout, p = 0.072; DMSO vs SR16584, p = 0.020; DMSO vs washout, p = 0.026; SR16584 vs washout, p = 0.020). *C*, The average total depolarization was also significantly decreased by application of SR16584 (repeated measures ANOVA with Tukey’s post hoc, *F*_*3,15*_ = 10.9; n = 6; control vs DMSO, p = 0.96; control vs SR16584, p = 0.0029; control vs washout, p = 0.32; DMSO vs SR16584, p = 0.007; DMSO vs washout, p = 0.25; SR16584 vs wash-out, p = 0.26).

### Repeated ACh pulses elicit temporal summation in VIP neurons

Thus far we have examined how isolated puffs of 1 mM ACh affected the excitability of VIP neurons. However, the time course and concentration of ACh released from cholinergic synapses onto VIP neurons in vivo is unknown. Previous studies show that the average firing rates of cholinergic PMT neurons in vivo tend to be rather low, typically less than a few Hz (Boucetta *et al*., 2014), but arousing sensory stimuli elicit brief bursts of firing that can reach 100 – 200 Hz (Reese *et al*., 1995*a*, 1995*b*; Sakai, 2012). In addition, our immunofluorescence data suggest that VIP neurons often receive multiple cholinergic inputs, which may reflect convergence from multiple PMT neurons. We therefore decided to test the effects of 10 Hz trains of ACh puffs, reasoning that VIP neurons would likely encounter this frequency of inputs in vivo. Based on the slow kinetics and limited desensitization of α_3_β_4_* nAChRs (David *et al*., 2010), we hypothesized that lower concentrations of ACh delivered in 10 Hz trains would elicit long-lasting excitation of VIP neurons due to temporal summation of cholinergic EPSPs. To test this, we made current-clamp recordings from VIP neurons while delivering trains of 1 – 10 puffs of 30 µM or 100 µM ACh (**Figure 7A,E**). We observed that as the number of puffs increased, VIP neurons increasingly depolarized and could transition from firing no spikes in response to a single ACh puff to firing trains of spikes in response multiple ACh puffs (**Figures 7B-D,F-H**). The amount of depolarization elicited by increasing numbers of ACh puffs, as measured by the average and normalized average area under the depolarization trace, produced input-output functions that were well described by linear fits, all of which had positive slopes (**Figure 7B,C,G,H**). Linear fits to the means of the normalized responses had slopes of 1.4 and 4.0, indicating that temporal summation was supralinear on average for both 30 µM and 100 µM puff trains, respectively (cyan data, **Figure 7D,H**). These results suggest that even if cholinergic synapses in vivo elicit smaller EPSPs, these EPSPs will be subject to temporal summation during periods of heightened PMT activity, thereby driving prolonged excitation of VIP neurons. This cholinergic excitation presumably combines with the ascending and local auditory inputs that VIP neurons receive (Goyer *et al*., 2019), reshaping auditory processing in VIP neurons and their postsynaptic targets.

**Figure 7:**
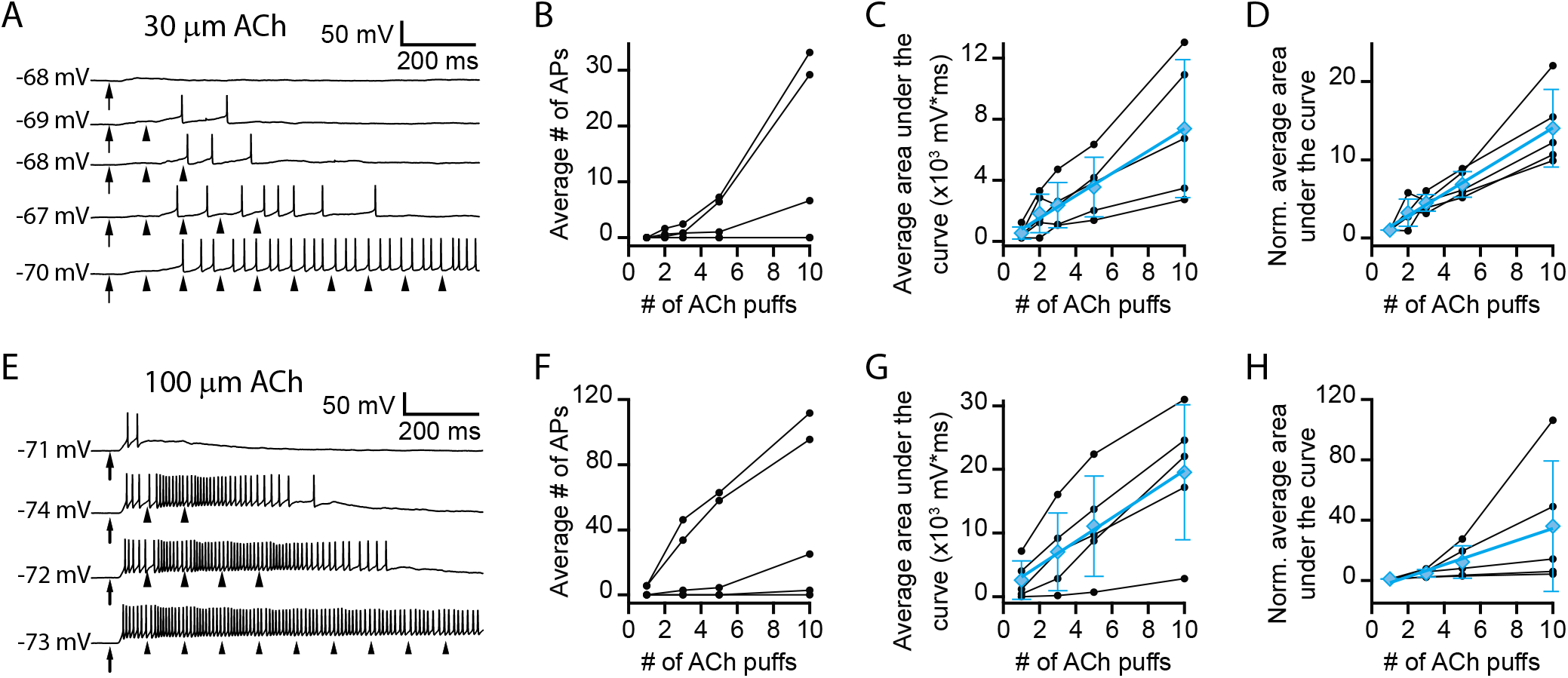
Trains of lower concentration ACh puffs elicit temporal summation and spiking in VIP neurons. *A*, Example traces show that 10 Hz trains of 10 ms, 30 µM ACh puffs elicited increased depolarization and probability of firing as the train duration increased from 1 – 10 puffs (top to bottom, respectively). *B*, In 3 out of 5 neurons, increasing the number of 30 µM ACh puffs led to spiking. For these 3 neurons, linear regressions yielded *r* values of 0.95, 0.97, and 0.97 and corresponding p values of 0.012, 0.006, 0.006. *C,D*, In 5 out of 5 neurons, increasing the number of 30 µM ACh puffs progressively increased the absolute (*C*) or normalized (*D*) total depolarization, measured as the area under the median-filtered curve, indicating that temporal summation occurred. Data for each neuron in *C,D* was well described by linear regression, with *r* values > 0.98 in each case. Cyan data in *C,D* show mean ± SD responses and corresponding linear fits (*C*, absolute data – slope = 730 mV*ms/puff; *r* = 0.99; n = 5; p = 0.002; *D*, normalized data – slope = 1.4 normalized units/puff; *r* = 0.99; n = 5; p = 0.0001). *E*, Example traces show that 10 Hz trains of 10 ms, 100 µM ACh puffs elicited increased depolarization and probability of firing as the train duration increased from 1 – 10 puffs (top to bottom, respectively). *F*, In 4 out of 5 neurons, increasing the number of 100 µM ACh puffs led to spiking. For 3 of these neurons, linear regressions yielded *r* values of 0.98, 0.99, and 0.96 and corresponding p values of 0.017, 0.013, and 0.037. *G,H*, In 5 out of 5 neurons, increasing the number of 100 µM ACh puffs progressively increased the absolute (*G*) or normalized (*H*) total depolarization, measured as the area under the median-filtered curve, indicating that temporal summation occurred. Data for each neuron in *G,H* was well described by linear regression, with *r* values > 0.97 in each case. Cyan data in *G,H* show mean ± SD responses and corresponding linear fits (absolute data – slope = 1860 mV*ms/puff; *r* = 0.99; n = 5; p = 0.002; normalized data – slope = 4.0 normalized units/puff; *r* = 0.99; n = 5; p = 0.011). In *A, E*, arrows and arrowheads indicate the times of ACh puffs, and voltages indicate resting membrane potential.

## Discussion

Here, we report the first cellular-level mechanism for cholinergic modulation in the auditory midbrain. Our data show that cholinergic terminals are routinely found in close proximity to the dendrites and somas of VIP neurons in the IC. In whole-cell recordings, brief applications of ACh to VIP neurons elicited surprisingly strong, long-lasting depolarizations and sustained inward currents. Despite the prolonged nature of these responses, they were not altered by blocking muscarinic receptors. Instead, using several nAChR antagonists, we determined that ACh excites VIP neurons mainly by activating α_3_β*_4_ nAChRs, with a small contribution from α_7_ nAChRs. α_3_β_4_* nAChRs are rare in the brain and have mainly been studied in the medial habenula and interpeduncular nucleus, where they play an important role in nicotine addiction (Sheffield *et al*., 2000; Grady *et al*., 2009; Scholze *et al*., 2012; Beiranvand *et al*., 2014). Our results uncover a novel role for α_3_β_4_* receptors in the central auditory pathway, revealing a potent neuromodulatory mechanism in which ACh can drive a sustained increase in the excitability of VIP neurons. Since VIP neurons project locally, to the auditory thalamus, and to several other auditory and non-auditory brain regions, cholinergic modulation of VIP neurons has the potential to exert widespread influence on auditory processing and its downstream effects.

### α_3_β_4_* nAChRs mediate prolonged depolarization of VIP neurons

Although nAChRs are often noted for driving fast and brief responses to cholinergic inputs, these effects are generally attributable to α_7_ nAChRs, which have fast kinetics and rapid desensitization (Christophe *et al*., 2002; Arroyo *et al*., 2012; Bennett *et al*., 2012). A growing number of studies have documented instances in which non-α_7_ nAChRs mediate longer-lasting changes in neuronal excitability. For example, in VIP interneurons in the auditory cortex, nicotine induces firing for up to several minutes, and this effect was blocked by DHβE, an α_4_β_2_* nAChR antagonist (Askew *et al*., 2019). In layer 1 of cerebral cortex, α_7_ nAChRs mediate an early, fast response to ACh, while non-α_7_ nAChRs mediate a later, slower response (Christophe *et al*., 2002; Arroyo *et al*., 2012; Bennett *et al*., 2012). Likewise, at the motoneuron-Renshaw cell synapse in the spinal cord, in combination with glutamatergic signaling, homomeric α_7_ nAChRs mediate an early, fast response to ACh, while α_4_β_2_* nAChRs mediate a slower, longer-lasting response (Lamotte d’Incamps & Ascher, 2008; d’Incamps *et al*., 2012; Lamotte d’Incamps *et al*., 2018).

While previous studies show that mRNA for α_7_, α_4_, and β_2_ nAChR subunits is common in the IC (Clarke *et al*., 1985; Wada *et al*., 1989; Morley & Happe, 2000; Happe & Morley, 2004; Bieszczad *et al*., 2012; Sottile *et al*., 2017), our data point to a limited role for α_7_ and no functional role for α_4_β_2_* nAChRs in VIP neurons. Instead, we found that α_3_β_4_* nAChRs are the dominant nAChRs in VIP neurons, mediating a strong, long-lasting depolarization in response to ACh application. Consistent with our observations, α_3_β_4_* nAChRs are capable of mediating sustained currents due to their slow desensitization, with time constants on the order of seconds, and relatively long single-channel open times and burst durations (David *et al*., 2010). In addition, in situ hybridization studies and binding studies have consistently shown that the IC is one of the few places in the brain where α_3_ and β_4_ nAChRs are expressed (Wada *et al*., 1989; Marks *et al*., 2002, 2006; Whiteaker *et al*., 2002; Salas *et al*., 2003; Gahring *et al*., 2004), and β_4_ knockout mice exhibit decreased α_3_ mRNA levels in the IC, supporting the hypothesis that α_3_ and β_4_ subunits interact in the IC (Salas *et al*., 2004). Our findings confirm and extend these results by providing the first evidence of a functional role for α_3_β_4_* nAChRs in the IC.

It is important to note that α_3_β_4_* nAChRs can have two stoichiometries, (α_3_β_4_)_2_α_3_ or (α_3_β_4_)_2_β_4_, and can also combine with a different fifth subunit, (α_3_β_4_)_2_X, where the fifth subunit can be α_2_, α_5_, α_6_, or β_3_ (Scholze & Huck, 2020). Previous studies indicate that the exact subunit composition of an α_3_β_4_* nAChR has some effect on Ca^2+^ permeability and desensitization rate, but generally little or no effect on the potency and efficacy of ACh (Wang *et al*., 1996; Gerzanich *et al*., 1998; Groot-Kormelink *et al*., 2001; Papke *et al*., 2010; Stokes & Papke, 2012). It will be important for future studies to determine which type or types of α_3_β_4_* nAChRs are expressed in VIP neurons.

### Trains of cholinergic inputs may drive long-lasting modulatory effects

To mimic more in vivo-like patterns of cholinergic input, we tested how VIP neurons responded to trains of ACh puffs. Our results show that, even at a relatively low application rate of 10 Hz, cholinergic EPSPs underwent substantial temporal summation in VIP neurons. This temporal summation allowed trains of 30 µM ACh puffs to transition from eliciting no spikes with 1 or 3 puffs to multiple spikes with 5 or 10 puffs. Trains of 100 µM ACh puffs elicited an even more pronounced increase in firing. Although ACh puffs probably do not match the concentration and time course of synaptically released ACh in vivo, our results show that ACh strongly excited VIP neurons under a range of conditions, uncovering a cellular mechanism that likely drives similar effects in vivo and highlighting the need for future experiments to build on these results.

It is well established that cholinergic PMT neurons, the source of cholinergic input to the IC, alter their firing rate as a function of behavioral state. For example, Boucetta et al. found that cholinergic PMT neurons fired maximally during the active wake and paradoxical sleep states (mean firing rates of 2.3 Hz and 3.7 Hz, respectively) and nearly ceased firing during slow wave sleep (0.04 Hz) (Boucetta *et al*., 2014). Sakai found similar changes across the sleep-wake cycle, but also found that arousing stimuli (a hand clap or an air puff) drove cholinergic PMT neurons to fire bursts of 2 – 5 spikes with instantaneous frequencies of 100 – 200 Hz (Sakai, 2012). Many PMT neurons also respond to sensory stimuli. For example, almost half of PMT neurons fire in response to auditory click stimuli, with half of these neurons firing short latency bursts (Reese *et al*., 1995*a*, 1995*b*). Such responses might be driven by the primary auditory cortex, which projects to the PMT (Schofield & Motts, 2009; Schofield, 2010). Furthermore, our immunofluorescence data suggest that VIP neurons often integrate multiple cholinergic inputs. It therefore seems likely that certain behavioral states and sensory stimuli drive cholinergic input to VIP neurons at rates sufficient to elicit the temporal summation we observed. Thus, our data support the hypothesis that cholinergic input from the PMT drives prolonged increases in the excitability in VIP neurons as a function of behavioral state and sensory input. Since VIP neurons project broadly within and beyond the IC, changes that alter cholinergic input to VIP neurons have the potential to drive wide-ranging changes in the excitability of the auditory and non-auditory circuits that VIP neurons target.

### Functional implications for auditory processing

Previous studies have identified clear roles for muscarinic signaling in the IC, including roles in cortically driven plasticity (Ji *et al*., 2001; Ji & Suga, 2009) and stimulus specific adaptation (Ayala & Malmierca, 2015), but the roles of nicotinic signaling in the IC are less clear. Psychophysical studies indicate that systemic nicotine exposure in non-smokers can enhance performance in auditory tasks (Harkrider & Hedrick, 2005; Knott *et al*., 2009; Pham *et al*., 2020). Intriguingly, recent work from Askew and colleagues suggests that systemic nicotine sharpens frequency tuning in the IC, which likely contributes to sharper tuning in auditory cortex and improved discrimination in behavioral tasks (Askew *et al*., 2017). However, the authors found that nicotine mainly suppressed activity in the IC. Since we found that activation of nAChRs increases VIP neuron excitability, this raises the interesting possibility that the local projections of VIP neurons might mainly target inhibitory neurons. Alternatively, a more complicated circuit interaction might occur. We plan to test how cholinergic modulation of VIP neurons shapes the auditory response properties of VIP neurons and their postsynaptic targets in future studies.

Finally, it is unknown how cholinergic signaling shapes the excitability of other IC neuron classes. The IC contains a rich diversity of neurons, but these have long proved difficult to reliably classify, making it difficult to investigate cholinergic modulation in a systematic way. Fortunately, in addition to VIP neurons, recent studies have identified GABAergic NPY neurons (Silveira *et al*., 2020) and glutamatergic CCK neurons (Kreeger *et al*., 2021) as distinct IC neuron classes. To gain a fuller understanding of how cholinergic modulation shapes auditory processing in the IC, it will be important to determine the diverse effects cholinergic modulation exerts on these and other, yet to be identified, neuron classes.

## Acknowledgements

We thank Pierre Apostolides, Don Caspary and Brett Schofield for discussions and helpful advice on this work, and Kevin Cruz-Colón for his help establishing the protocol for the immunohistochemistry experiments. This work was supported by a Rackham Graduate Student Research Grant (University of Michigan) and National Institutes of Health grants T32 DC005356 (Gabriel Corfas), F31 DC019292 (LMR), and R01 DC018284 (MTR).

## References

Anand R, Nelson ME, Gerzanich V, Wells GB & Lindstrom J (1998). Determinants of Channel Gating Located in the N-Terminal Extracellular Domain of Nicotinic α7 Receptor. J Pharmacol Exp Ther 287, 469–479.

Arroyo S, Bennett C, Aziz D, Brown SP & Hestrin S (2012). Prolonged Disynaptic Inhibition in the Cortex Mediated by Slow, Non-α7 Nicotinic Excitation of a Specific Subset of Cortical Interneurons. J Neurosci 32, 3859–3864.

Askew C, Intskirveli I & Metherate R (2017). Systemic Nicotine Increases Gain and Narrows Receptive Fields in A1 via Integrated Cortical and Subcortical Actions. eNeuro; DOI: 10.1523/ENEURO.0192-17.2017.

Askew CE, Lopez AJ, Wood MA & Metherate R (2019). Nicotine excites VIP interneurons to disinhibit pyramidal neurons in auditory cortex. Synapsee22116.

Ayala YA & Malmierca MS (2015). Cholinergic Modulation of Stimulus-Specific Adaptation in the Inferior Colliculus. Journal of Neuroscience 35, 12261–12272.

Beiranvand F, Zlabinger C, Orr-Urtreger A, Ristl R, Huck S & Scholze P (2014). Nicotinic acetylcholine receptors control acetylcholine and noradrenaline release in the rodent habenulo-interpeduncular complex. Br J Pharmacol 171, 5209–5224.

Bennett C, Arroyo S, Berns D & Hestrin S (2012). Mechanisms Generating Dual-Component Nicotinic EPSCs in Cortical Interneurons. J Neurosci 32, 17287–17296.

Bieszczad KM, Kant R, Constantinescu CC, Pandey SK, Kawai HD, Metherate R, Weinberger NM & Mukherjee J (2012). Nicotinic acetylcholine receptors in rat forebrain that bind 18F-nifene: relating PET imaging, autoradiography and behavior. Synapse 66, 418–434.

Boucetta S, Cissé Y, Mainville L, Morales M & Jones BE (2014). Discharge Profiles across the Sleep– Waking Cycle of Identified Cholinergic, GABAergic, and Glutamatergic Neurons in the Pontomesencephalic Tegmentum of the Rat. J Neurosci 34, 4708–4727.

Castro NG & Albuquerque EX (1993). Brief-lifetime, fast-inactivating ion channels account for the α-bungarotoxin-sensitive nicotinic response in hippocampal neurons. Neuroscience Letters 164, 137–140.

Christophe E, Roebuck A, Staiger JF, Lavery DJ, Charpak S & Audinat E (2002). Two Types of Nicotinic Receptors Mediate an Excitation of Neocortical Layer I Interneurons. Journal of Neurophysiology 88, 1318–1327.

Clarke P, Schwartz R, Paul S, Pert C & Pert A (1985). Nicotinic binding in rat brain: autoradiographic comparison of [3H]acetylcholine, [3H]nicotine, and [125I]-alpha-bungarotoxin. J Neurosci 5, 1307–1315.

Dani JA & Bertrand D (2007). Nicotinic acetylcholine receptors and nicotinic cholinergic mechanisms of the central nervous system. Annu Rev Pharmacol Toxicol 47, 699–729.

David R, Ciuraszkiewicz A, Simeone X, Orr-Urtreger A, Papke RL, McIntosh JM, Huck S & Scholze P (2010). Biochemical and functional properties of distinct nicotinic acetylcholine receptors in the superior cervical ganglion of mice with targeted deletions of nAChR subunit genes. Eur J Neurosci 31, 978–993.

Dodge FA & Rahamimoff R (1967). Co-operative action of calcium ions in transmitter release at the neuromuscular junction. J Physiol 193, 419–432.

Farley GR, Morley BJ, Javel E & Gorga MP (1983). Single-unit responses to cholinergic agents in the rat inferior colliculus. Hearing Research 11, 73–91.

Felix RA, Chavez VA, Novicio DM, Morley BJ & Portfors CV (2019). Nicotinic acetylcholine receptor subunit α 7 -knockout mice exhibit degraded auditory temporal processing. Journal of Neurophysiology 122, 451–465.

Forman CJ, Tomes H, Mbobo B, Burman RJ, Jacobs M, Baden T & Raimondo JV (2017). Openspritzer: an open hardware pressure ejection system for reliably delivering picolitre volumes. Scientific Reports 7, 2188.

Gahring LC, Persiyanov K & Rogers SW (2004). Neuronal and astrocyte expression of nicotinic receptor subunit β4 in the adult mouse brain. Journal of Comparative Neurology 468, 322–333.

Gerzanich V, Wang F, Kuryatov A & Lindstrom J (1998). alpha 5 Subunit alters desensitization, pharmacology, Ca++ permeability and Ca++ modulation of human neuronal alpha 3 nicotinic receptors. J Pharmacol Exp Ther 286, 311–320.

Gillet C, Goyer D, Kurth S, Griebel H & Kuenzel T (2018). Cholinergic innervation of principal neurons in the cochlear nucleus of the Mongolian gerbil. J Comp Neurol 526, 1647–1661.

Goyer D, Kurth S, Gillet C, Keine C, Rübsamen R & Kuenzel T (2016). Slow Cholinergic Modulation of Spike Probability in Ultra-Fast Time-Coding Sensory Neurons. eNeuro; DOI: 10.1523/ENEURO.0186-16.2016.

Goyer D, Silveira MA, George AP, Beebe NL, Edelbrock RM, Malinski PT, Schofield BR & Roberts MT (2019). A novel class of inferior colliculus principal neurons labeled in vasoactive intestinal peptide-Cre mice. eLife 8, e43770.

Grady SR, Moretti M, Zoli M, Marks MJ, Zanardi A, Pucci L, Clementi F & Gotti C (2009). Rodent Habenulo–Interpeduncular Pathway Expresses a Large Variety of Uncommon nAChR Subtypes, But Only the α3β4 and α3β3β4 Subtypes Mediate Acetylcholine Release. J Neurosci 29, 2272– 2282.

Groot-Kormelink PJ, Boorman JP & Sivilotti LG (2001). Formation of functional α3β4α5 human neuronal nicotinic receptors in Xenopus oocytes: a reporter mutation approach. Br J Pharmacol 134, 789– 796.

Habbicht H & Vater M (1996). A microiontophoretic study of acetylcholine effects in the inferior colliculus of horseshoe bats: implications for a modulatory role. Brain Research 724, 169–179.

Happe HK & Morley BJ (2004). Distribution and postnatal development of α7 nicotinic acetylcholine receptors in the rodent lower auditory brainstem. Developmental Brain Research 153, 29–37.

Harkrider AW & Hedrick MS (2005). Acute effect of nicotine on auditory gating in smokers and non-smokers. Hearing Research 202, 114–128.

d’Incamps BL, Krejci E & Ascher P (2012). Mechanisms Shaping the Slow Nicotinic Synaptic Current at the Motoneuron–Renshaw Cell Synapse. J Neurosci 32, 8413–8423.

Ji W, Gao E & Suga N (2001). Effects of Acetylcholine and Atropine on Plasticity of Central Auditory Neurons Caused by Conditioning in Bats. Journal of Neurophysiology 86, 211–225.

Ji W & Suga N (2009). Tone-Specific and Nonspecific Plasticity of Inferior Colliculus Elicited by Pseudo-Conditioning: Role of Acetylcholine and Auditory and Somatosensory Cortices. Journal of Neurophysiology 102, 941–952.

Jones BE (1991). Paradoxical sleep and its chemical/structural substrates in the brain. Neuroscience 40, 637–656.

Knott V, Shah D, Millar A, McIntosh J, Fisher D, Blais C & Ilivitsky V (2012). Nicotine, Auditory Sensory Memory, and sustained Attention in a Human Ketamine Model of Schizophrenia: Moderating Influence of a Hallucinatory Trait. Front Pharmacol 3, 172.

Knott VJ, Bolton K, Heenan A, Shah D, Fisher DJ & Villeneuve C (2009). Effects of acute nicotine on event-related potential and performance indices of auditory distraction in nonsmokers. Nicotine & Tobacco Research 11, 519–530.

Kozak R, Bowman EM, Latimer MP, Rostron CL & Winn P (2005). Excitotoxic lesions of the pedunculopontine tegmental nucleus in rats impair performance on a test of sustained attention. Exp Brain Res 162, 257–264.

Kreeger LJ, Connelly CJ, Mehta P, Zemelman BV & Golding NL (2021). Excitatory cholecystokinin neurons of the midbrain integrate diverse temporal responses and drive auditory thalamic subdomains. Proc Natl Acad Sci U S A; DOI: 10.1073/pnas.2007724118.

Lamotte d’Incamps B & Ascher P (2008). Four Excitatory Postsynaptic Ionotropic Receptors Coactivated at the Motoneuron–Renshaw Cell Synapse. J Neurosci 28, 14121–14131.

Lamotte d’Incamps B, Zorbaz T, Dingova D, Krejci E & Ascher P (2018). Stoichiometry of the Heteromeric Nicotinic Receptors of the Renshaw Cell. J Neurosci 38, 4943–4956.

Madisen L, Zwingman TA, Sunkin SM, Oh SW, Zariwala HA, Gu H, Ng LL, Palmiter RD, Hawrylycz MJ, Jones AR, Lein ES & Zeng H (2010). A robust and high-throughput Cre reporting and characterization system for the whole mouse brain. Nat Neurosci 13, 133–140.

Marks MJ, Whiteaker P & Collins AC (2006). Deletion of the α7, β2, or β4 Nicotinic Receptor Subunit Genes Identifies Highly Expressed Subtypes with Relatively Low Affinity for [3H]Epibatidine. Mol Pharmacol 70, 947–959.

Marks MJ, Whiteaker P, Grady SR, Picciotto MR, McIntosh JM & Collins AC (2002). Characterization of [125I]epibatidine binding and nicotinic agonist-mediated 86Rb+ efflux in interpeduncular nucleus and inferior colliculus of β2 null mutant mice. Journal of Neurochemistry 81, 1102–1115.

Mike A, Castro NG & Albuquerque EX (2000). Choline and acetylcholine have similar kinetic properties of activation and desensitization on the α7 nicotinic receptors in rat hippocampal neurons. Brain Research 882, 155–168.

Millar NS & Gotti C (2009). Diversity of vertebrate nicotinic acetylcholine receptors. Neuropharmacology 56, 237–246.

Morley BJ & Happe HK (2000). Cholinergic receptors: dual roles in transduction and plasticity. Hearing Research 147, 104–112.

Motts SD & Schofield BR (2009). Sources of cholinergic input to the inferior colliculus. Neuroscience 160, 103–114.

Noftz WA, Beebe NL, Mellott JG & Schofield BR (2020). Cholinergic Projections From the Pedunculopontine Tegmental Nucleus Contact Excitatory and Inhibitory Neurons in the Inferior Colliculus. Front Neural Circuits; DOI: 10.3389/fncir.2020.00043.

Papke RL, Dwoskin LP, Crooks PA, Zheng G, Zhang Z, McIntosh JM & Stokes C (2008). Extending the analysis of nicotinic receptor antagonists with the study of alpha6 nicotinic receptor subunit chimeras. Neuropharmacology 54, 1189–1200.

Papke RL, Meyer E, Nutter T & Uteshev VV (2000). α7 Receptor-selective agonists and modes of α7 receptor activation. European Journal of Pharmacology 393, 179–195.

Papke RL, Sanberg PR & Shytle RD (2001). Analysis of Mecamylamine Stereoisomers on Human Nicotinic Receptor Subtypes. J Pharmacol Exp Ther 297, 646–656.

Papke RL, Wecker L & Stitzel JA (2010). Activation and Inhibition of Mouse Muscle and Neuronal Nicotinic Acetylcholine Receptors Expressed in Xenopus Oocytes. J Pharmacol Exp Ther 333, 501–518.

Pham CQ, Kapolowicz MR, Metherate R & Zeng F-G (2020). Nicotine enhances auditory processing in healthy and normal-hearing young adult nonsmokers. Psychopharmacology (Berl) 237, 833–840.

Rees CL, Moradi K & Ascoli GA (2017). Weighing the Evidence in Peters’ Rule: Does Neuronal Morphology Predict Connectivity? Trends Neurosci 40, 63–71.

Reese NB, Garcia-Rill E & Skinner RD (1995a). Auditory input to the pedunculopontine nucleus: I. Evoked potentials. Brain Research Bulletin 37, 257–264.

Reese NB, Garcia-Rill E & Skinner RD (1995b). Auditory input to the pedunculopontine nucleus: II. Unit responses. Brain Research Bulletin 37, 265–273.

Sakai K (2012). Discharge properties of presumed cholinergic and noncholinergic laterodorsal tegmental neurons related to cortical activation in non-anesthetized mice. Neuroscience 224, 172–190.

Salas R, Cook KD, Bassetto L & De Biasi M (2004). The α3 and β4 nicotinic acetylcholine receptor subunits are necessary for nicotine-induced seizures and hypolocomotion in mice. Neuropharmacology 47, 401–407.

Salas R, Pieri F, Fung B, Dani JA & De Biasi M (2003). Altered Anxiety-Related Responses in Mutant Mice Lacking the β4 Subunit of the Nicotinic Receptor. J Neurosci 23, 6255–6263.

Schofield BR (2010). Projections from auditory cortex to midbrain cholinergic neurons that project to the inferior colliculus. Neuroscience 166, 231.

Schofield BR & Motts SD (2009). Projections from auditory cortex to cholinergic cells in the midbrain tegmentum of guinea pigs. Brain Res Bull 80, 163–170.

Schofield BR, Motts SD & Mellott JG (2011). Cholinergic Cells of the Pontomesencephalic Tegmentum: Connections with Auditory Structures from Cochlear Nucleus to Cortex. Hear Res 279, 85–95.

Scholze P & Huck S (2020). The α5 Nicotinic Acetylcholine Receptor Subunit Differentially Modulates α4β2* and α3β4* Receptors. Front Synaptic Neurosci; DOI: 10.3389/fnsyn.2020.607959.

Scholze P, Koth G, Orr-Urtreger A & Huck S (2012). Subunit composition of α5-containing nicotinic receptors in the rodent habenula. J Neurochem 121, 551–560.

Sheffield EB, Quick MW & Lester RAJ (2000). Nicotinic acetylcholine receptor subunit mRNA expression and channel function in medial habenula neurons. Neuropharmacology 39, 2591–2603.

Silveira MA, Anair JD, Beebe NL, Mirjalili P, Schofield BR & Roberts MT (2020). Neuropeptide Y Expression Defines a Novel Class of GABAergic Projection Neuron in the Inferior Colliculus. J Neurosci 40, 4685–4699.

Smucny J, Olincy A, Rojas DC & Tregellas JR (2016). Neuronal effects of nicotine during auditory selective attention in schizophrenia. Hum Brain Mapp 37, 410–421.

Sottile SY, Hackett TA, Cai R, Ling L, Llano DA & Caspary DM (2017). Presynaptic Neuronal Nicotinic Receptors Differentially Shape Select Inputs to Auditory Thalamus and Are Negatively Impacted by Aging. J Neurosci 37, 11377–11389.

Stokes C & Papke RL (2012). Use of an α3-β4 nicotinic acetylcholine receptor subunit concatamer to characterize ganglionic receptor subtypes with specific subunit composition reveals species-specific pharmacologic properties. Neuropharmacology 63, 538–546.

Taniguchi H, He M, Wu P, Kim S, Paik R, Sugino K, Kvitsiani D, Kvitsani D, Fu Y, Lu J, Lin Y, Miyoshi G, Shima Y, Fishell G, Nelson SB & Huang ZJ (2011). A resource of Cre driver lines for genetic targeting of GABAergic neurons in cerebral cortex. Neuron 71, 995–1013.

Wada E, Wada K, Boulter J, Deneris E, Heinemann S, Patrick J & Swanson LW (1989). Distribution of alpha 2, alpha 3, alpha 4, and beta 2 neuronal nicotinic receptor subunit mRNAs in the central nervous system: a hybridization histochemical study in the rat. J Comp Neurol 284, 314–335.

Wang F, Gerzanich V, Wells GB, Anand R, Peng X, Keyser K & Lindstrom J (1996). Assembly of Human Neuronal Nicotinic Receptor α5 Subunits with α3, β2, and β4 Subunits *. Journal of Biological Chemistry 271, 17656–17665.

Whiteaker P, Peterson CG, Xu W, McIntosh JM, Paylor R, Beaudet AL, Collins AC & Marks MJ (2002). Involvement of the α3 Subunit in Central Nicotinic Binding Populations. J Neurosci 22, 2522– 2529.

Zaveri N, Jiang F, Olsen C, Polgar W & Toll L (2010). Novel alpha3beta4 nicotinic acetylcholine receptor-selective ligands. Discovery, structure-activity studies and pharmacological evaluation. J Med Chem 53, 8187–8191.

Zhang C, Beebe NL, Schofield BR, Pecka M & Burger RM (2021). Endogenous Cholinergic Signaling Modulates Sound-Evoked Responses of the Medial Nucleus of the Trapezoid Body. J Neurosci 41, 674–688.

Zhang L, Wu C, Martel DT, West M, Sutton MA & Shore SE (2019). Remodeling of cholinergic input to the hippocampus after noise exposure and tinnitus induction in Guinea pigs. Hippocampus 29, 669– 682.

Zheng QY, Johnson KR & Erway LC (1999). Assessment of hearing in 80 inbred strains of mice by ABR threshold analyses. Hear Res 130, 94–107.

